# Extraction of common task features in EEG-fMRI data using coupled tensor-tensor decomposition

**DOI:** 10.1101/685941

**Authors:** Yaqub Jonmohamadi, Suresh Muthukumaraswamy, Joseph Chen, Jonathan Roberts, Ross Crawford, Ajay Pandey

## Abstract

The fusion of simultaneously recorded EEG and fMRI data is of great value to neuroscience research due to the complementary properties of the individual modalities. Traditionally, techniques such as PCA and ICA, which rely on strong strong non-physiological assumptions such as orthogonality and statistical independence, have been used for this purpose. Recently, tensor decomposition techniques such as parallel factor analysis have gained more popularity in neuroimaging applications as they are able to inherently contain the multidimensionality of neuroimaging data and achieve uniqueness in decomposition without imposing strong assumptions. Previously, the coupled matrix-tensor decomposition (CMTD) has been applied for the fusion of the EEG and fMRI. Only recently the coupled tensor-tensor decomposition (CTTD) has been proposed. Here for the first time, we propose the use of CTTD of a 4th order EEG tensor (space, time, frequency, and participant) and 3rd order fMRI tensor (space, time, participant), coupled partially in time and participant domains, for the extraction of the task related features in both modalities. We used both the sensor-level and source-level EEG for the coupling. The phase shifted paradigm signals were incorporated as the temporal initializers of the CTTD to extract the task related features. The validation of the approach is demonstrated on simultaneous EEG-fMRI recordings from six participants performing an N-Back memory task. The EEG and fMRI tensors were coupled in 9 components out of which 7 components had a high correlation (more than 0.85) with the task. The result of the fusion recapitulates the well-known attention network as being positively, and the default mode network working negatively time-locked to the memory task.

## 1 Introduction

Functional magnetic resonance images (fMRI) (Ogawa et al. 1990) measures the slowly changing blood oxygenation level dependent (BOLD) signals as an indirect measure of neural activity and is a popular neuroimaging technique thanks to its high spatial resolution (typically a few millimeters). On the other hand, electroencephalography (EEG) measures neural activity, almost directly, using scalp electrodes. Despite the superior temporal resolution (millisecond), the spatial reconstruction of EEG data is an ill-posed problem (projection of the 2D scalp measurement into the 3D brain space) and has a poor spatial resolution. Hence, the fusion/integration of these two modalities is of great value due to their complementary spatiotemporal properties (Babiloni et al. 2004).

Approaches to the fusion of EEG and fMRI are divided into two main categories: model-driven and data-driven. In model-driven methods, one modality is estimated from the other using computational biophysical models. Since precise knowledge about the neuronal substrates is rarely available, the use of model-driven methods has been decreasing (Valdes-Sosa et al. 2009; Ferdowsi et al. 2015). On the other hand, data-driven methods incorporate both modalities to estimate fused components. There is an extensive literature on the use of data-driven approaches for EEG-fMRI integration which are mostly based on decomposing the EEG and fMRI into different components. Traditional matrix decomposition techniques such as principal component analysis (PCA) and independent component analysis (ICA) have been used as the primary engines for the preprocessing, feature extraction, and in a more versatile way, the fusion of the two neuroimaging techniques. One reason for the popularity of these two techniques is the convenience of presenting the time-varying EEG and fMRI as a matrix of time × space (channel or voxels).

PCA has a long history of application in EEG preprocessing such as in artifact rejection and extraction of meaningful brain activity (Lagerlund et al. 1997; Soong and Koles 1995). A basic problem is that components are defined by only two signatures (space and time) which are not determined uniquely, therefore orthogonality is imposed between the corresponding signatures of different components (Miwakeichi et al. 2004). ICA has been used extensively in both EEG and fMRI literature (Makeig et al. 2004; Jonmohamadi et al. 2014a; Jonmohamadi and Jones 2015) and is a popular tool for space/time decomposition. While it avoids the orthogonality constraint, uniqueness, however, is achieved at the price of imposing an even stronger non-physiological constraint, namely, that sources are statistically independent (Miwakeichi et al. 2004).

In a typical EEG or fMRI experiment the data has higher dimension than time×space and it could go up to seven dimensions (Cong et al. 2015), e.g., in case of EEG time×space×frequency×trial×condition× participant×group. In order to use ICA and/or PCA for these higher dimensional data, unfolding of some modalities onto others and reducing the data dimension into a matrix is necessary, which is typically done by concatenation or stacking of the data (Delorme and Makeig 2004; Eichele et al. 2011; Dien 2012; Calhoun and Adali 2012; Cong et al. 2013). Such unfolding inevitably loses some potentially existing interactions between/among the folded modes (Cong et al. 2015) and makes the interpretation of the results difficult (Mrup et al. 2006).

Considering that the EEG and fMRI data can be expressed conveniently as a three or higher dimensional array (tensor) it is possible and favorable to use tensor decomposition techniques for the purpose of breaking the data into components with corresponding signatures from each dimension. One such techniques is the parallel factor analysis (PARAFAC), also known as canonical polyadic decomposition, (Carroll and Chang 1970; Harshman 1970). Besides being able to conveniently deal with multidimensional data, the second main advantage of the PARAFAC over the PCA and ICA is that uniqueness is achieved without the need for imposing strong physiologically irrelevant assumptions such as orthogonality and statistical independence.

### 1.1 Tensor factorization-based EEG-fMRI analysis

Tensors are higher order generalization of matrices, i.e., multiway arrays, which can represent additional types of variables in their higher dimensions (Hunyadi et al. 2016). Recently, tensor decomposition techniques such as PARAFAC or Tucker have become attractive in signal processing (Cichocki et al. 2015; Cong et al. 2015; Sen and Parhi 2017) and in neuroimaging applications for being able to naturally present the inherently multidimensional data and preserve their structural information defined by inter-dependencies among various modes of variability such as time, space, participant, or frequency (Hunyadi et al. 2016). In neuroimaging, often one or more sources of variability exist between the measurement from different modalities, for example, neural activity might have a similar temporal pattern in EEG and MEG (magnetoencephalography) (Hunyadi et al. 2016). Alternatively, using mathematical manipulations, such as convolution of the EEG (or MEG) signals/features with a HRF, it can be assumed that EEG and fMRI share temporal similarities. Other sources of variability could include participant, task, and group.

So far, all except one tensor-based fusion of EEG-fMRI have been based on different variants of the coupled matrix-tensor factorization (CMTF) (Acar et al. 2013), in which data is structured in such a way that a matrix contains fMRI data and a 3rd order tensor contains the EEG activity and it is assumed that there is one common mode of variability between the matrix and the tensor, for example, time (Martinez-Montes et al. 2004) or participant (Acar et al. 2013, 2014; Hunyadi et al. 2016, 2017). The factorization of the structured data is achieved by imposing constraints on the optimization algorithm. Hence, the CMTF is based on the strong assumption that components in the shared dimension are equal. To relax this assumption, several alternatives have been introduced, such as Advanced CMTF (Karahan et al. 2015; Acar et al. 2014) which allows both shared and nonshared components in the common mode between the matrix and the tensor or Relaxed Advanced CMTF (Rivet et al. 2015), soft coupling (Seichepine et al. 2014), and approximate coupling (Farias et al. 2016) which provides similarity rather than the equivalence between the common components. Only recently, Chatzichristos et al. (Chatzichristos et al. 2018) for the first time, used the coupled tensor-tensor decomposition (CTTD) of the EEG and fMRI data fusion. They also used the so called ‘soft’ coupling approach to alleviate the strong assumption related to the canonical haemodynamic response function (HRF). They demonstrated their superiority over the ICA based fusion approaches (Calhoun et al. 2009, 2006; Mijovi et al. 2012) using simulated data.

To the best knowledge of the authors only two papers have reported the use of source-level EEG for fusion with fMRI data (Karahan et al. 2015; Jonmohamadi et al. 2018) and all the other mentioned EEG-fMRI technique have used sensor-level EEG, and to estimate the brain maps, post-processing was applied to the fused sensor-level EEG maps. The source-level EEG has two main advantages over the sensor-level counterpart: firstly, data is less mixed as it is spatially filtered using typically a minimum norm or a minimum variance filter. Secondly, the fused components are accompanied directly with the EEG brain maps and it is a desirable property to be able to compare the fMRI and EEG brain maps for the same activity directly. Since we are using both sensor and source-level EEG from multiple bands, the result of the fused components will be accompanied with scalp and brain maps from each frequency band as well as the corresponding brain map from the fMRI.

## 2 Methods

In this manuscript, the *italic* lower case refers to scalar (*a*), ***bold italic*** lower case refers to vectors (***a***), ***bold italic*** upper case refers to matrices (***A***), and calligraphic upper case letters refers to tensors (𝒜). <u>The glossary of the mathematical characters used in the following sections is provided in</u> table 1.

**Table 1:**
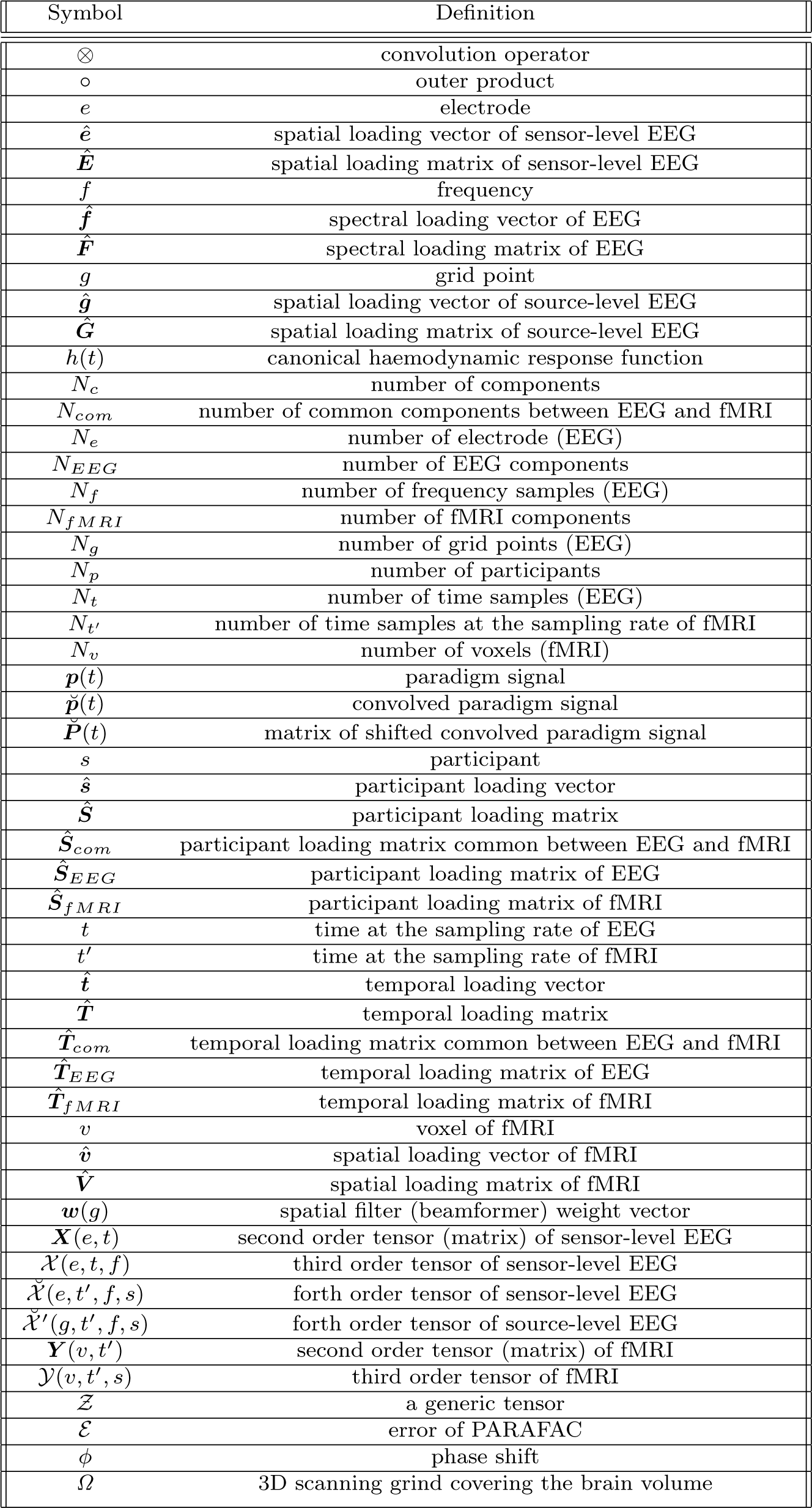
Glossary of the characters and operands used in the manuscript.

### 2.1 Data structure

The millisecond resolution of the EEG provides rich temporal information on the dynamic changes of brain activities. However, since EEG captures activity from large number of physiological and non-physiological sources using a limited number of sensors, mathematical operations are required to filter noise sources and at the same time extract the spectral, temporal, and spatial information of the sources of interest. By default the recorded EEG from *N*_*e*_ electrodes, during *N*_*t*_ time samples, i.e., 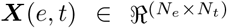, does not contain the spectral information as a 3rd dimension. Hence time-frequency analysis techniques such as wavelet (e.g., (Kronland-Martinet et al. 1987)) can be applied to extract the spectral signatures of the activities at each electrode, or alternatively, temporal band-pass filtering can be applied to divide the wide band EEG, **X**(*e, t*), into the popular delta (0-4 Hz), theta (4-8 Hz), alpha (8-12 Hz), beta (12-28 Hz), and gamma (+28 Hz) sub-bands,

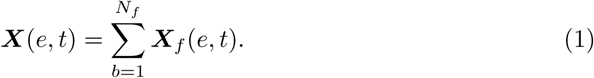

Using the band-pass filtering the EEG data can be presented in a 3rd order tensor

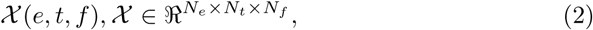

where *f* refers to frequency and *N*_*f*_ is the number of the sub-bands.

One inherent limitation of EEG is that the source-level activity (3D brain space) is recorded using the sensor-level electrodes (2D space). As a result, the activities of different sources are highly overlapped at the sensor-level. Inverse solutions such as minimum-variance (Van Veen et al. 1997; Jonmohamadi et al. 2014b) spatial filters can be used to project the sensor-level data into the source-level *Ω*, to reduce the overlap of the sources and at the same time estimate the brain maps associated with certain activities. A scanning grid is required to cover the brain space and spatial filter coefficients ***w***(*g*) should be applied for every point of the scanning grid *g* to estimate the EEG activity from that point,

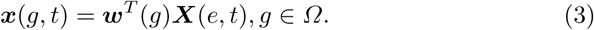

The projected EEG into the brain space is a matrix with *N*_*g*_ grid points and *N*_*t*_ time points 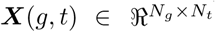, which has a substantial increase in the space domain from *N*_*e*_ to *N*_*g*_, compared with the sensor-level EEG, for example, 64 sensors compared with a few thousands voxels. Similar to the sensor-level EEG, the source-level EEG can also be band-pass filtered (***X***_*f*_ (*g, t*)) to create the 3rd order tensor,

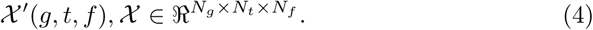

The fMRI data from *N*_*v*_ voxels and *N*_*t*_*I* time points, i.e., 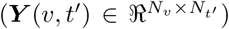, is recorded with a much slower sampling rate (typically a few seconds) compared with the EEG. In order to temporally relate the EEG with the fMRI, the preprocessed EEG data is convolved with a canonical HRF (Glover 1999) function to take the haemodynamic delay into account,

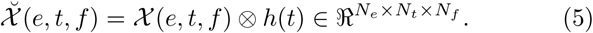

Since the participants are performing a task, the experimentally specified paradigm signal could be used as a temporal constraint, to extract the task and non-task related EEG and fMRI intervals. In our example data the paradigm ***p***(*t*) (often a boxcar in fMRI studies) refers to a 30 s of working-memory task followed by a 30 s of non-task (resting state) period, which occurs for 9 cycles. The paradigm signal is also convolved with the canonical HRF.

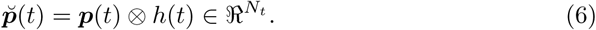

It is known that the brain has a lag structure (Mitra et al. 2014; Feige et al. 2017), meaning different brain regions have different HRF. To account for this varying HRFs, the convolved paradigm signal was shifted several times to create a matrix of the paradigm signals with each row of it having slightly a shift in phase compared with the previous row. This matrix could be used as a temporal constraint to extract the corresponding spatial, spectral and participant related features

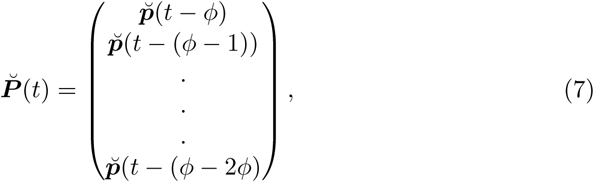

where *ϕ* is the number of the time sample shifts. Both the EEG and paradigm signals are downsampled to the fMRI sampling rate of 0.4545 Hz (*t* ⇒ *t*′).

Another dimension of data arises when the recording involves several participants (*N*_*s*_) which is typical in EEG and fMRI studies. Therefore, the fMRI is presented by a 3rd order tensor and the sensor-level or source-level EEG with a 4th order tensor:

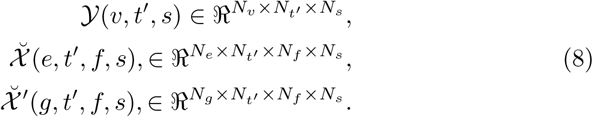

### 2.2 PARAFAC

PARAFAC approximates the original tensor, as the sum of *N*_*c*_ rank one tensors,

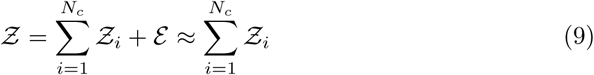

Where *ε* is the error tensor, and *Ƶ*_*i*_ is rank 1 tensor corresponding to component *i*. In the case of EEG and fMRI this can be written as

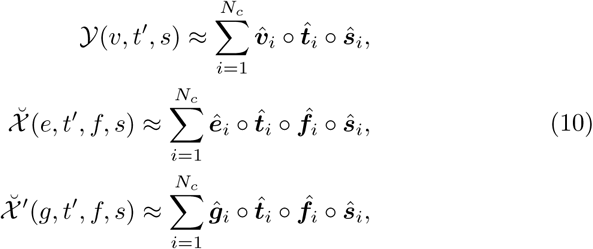

where ∘ is the outer product and 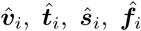, and 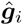 are known as “loading vectors” which correspond to the spatial, temporal and participants’ signatures of the *i*th component, in this work. Here for the simplicity the loading vectors are also referred to as ‘components’. Similarly, the loading matrices 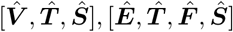, and 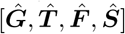 contain the spatial, temporal, spectral, and participants signatures of all components for fMRI and EEG data.

### 2.3 Coupled tensor-tensor factorization

In the case of coupled tensor-tensor factorization for the fMRI and sensor-level EEG, the original CMTF (Sorber et al. 2015) can be written as

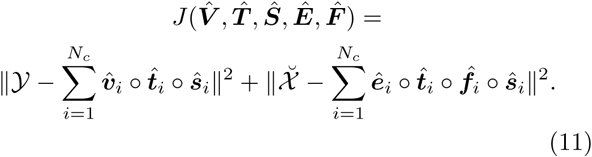

In the above cost function, the fMRI and EEG are coupled in time and participant modes. The coupled components are exactly the same in the two tensor data which is known to be a strong assumption. Moreover, it is highly likely that the EEG and fMRI have different data ranks, i.e., *N*_*EEG*_ and *N*_*fMRI*_. One way to alleviate this assumption is to use partial coupling, where only some of the components in time and participants are coupled between the two tensors

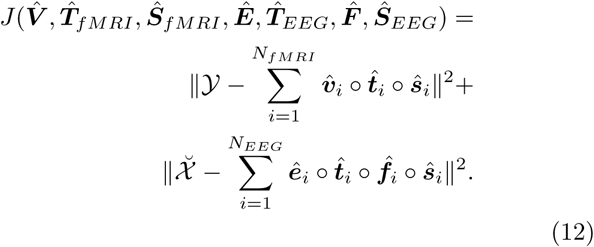

The 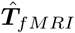 and 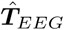 are partially the same which is also the case with ***Ŝ***_*fMRI*_ and ***Ŝ***_*EEG*_:

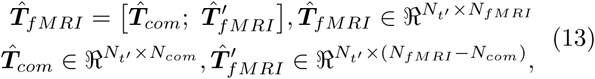

and similarly

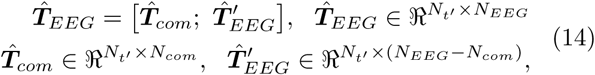

where *N*_*com*_ is the number of the common components in time and participant domain between EEG and fMRI and ‘;’ indicates vertically concatenated matrices. Similar equations to 13 and 14 are applicable for the participant loading matrices (***Ŝ***_*f MRI*_ and ***Ŝ***_*EEG*_).

Since the subjects are performing a common task, i.e., 30 s of 2-Back followed by 30 s of 0-Back memory tasks, the temporal signatures of desired EEG and fMRI features are approximately known

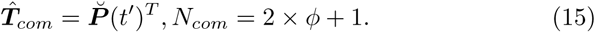

Therefore, the 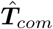 can be used as a temporal constraint for solving Eg. (12) and its corresponding spatial, spectral, and participant loading matrices can be extracted as task related features. However, applying the 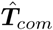 as the partially known temporal loading matrix is also a strong assumption and fluctuations in the power of the EEG and fMRI are expected which will not match with the shape of the 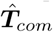. In the case of the coupled data factorization, the initialization of the loading matrices is known to be an important step (Vervliet et al. 2016). Hence, rather than considering the 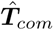 as a known loading matrix, it is was used as the initializer of the coupled EEG and fMRI loading matrix.

There are several methods on deciding the number of the components (*N*_*EEG*_ and *N*_*fMRI*_). However, these methods could show substantially different results. For example, the rank estimate provided by TensorLab toolbox (Vervliet et al. 2016) indicates 44 is the number of components related to the sensor-level EEG which has a low error in PARAFAC estimation, whereas, the core consistency diagnosis of N-way toolbox (Andersson and Bro 2000) indicated 92% score when only *N*_*EEG*_ = 5. While setting *N*_*EEG*_ = 44 resulted in many similar/identical components, the *N*_*EEG*_ = 5 appeared to be too small. The trial and error was suggested as alternative in (Vervliet et al. 2016). Using the trial- and error approach it appeared setting *N*_*EEG*_ = 20 and *N*_*fMRI*_ = 14 results in components which are not similar/identical and the same time most of them are reproducible through rerunning the PARAFAC. Similarly, the *N*_*com*_ was set to 9, which covered +4.4 s to −4.4 s with 1.1 s steps with respect to the paradigm signal. There were 7 coupled components reproducible through rerunning the coupled tensor-tensor factorization. The coupling of the EEG and fMRI could be done in two ways, source-level EEG and fMRI or sensor-level EEG and fMRI. They result in similar features however, the earlier is more time consuming due to the size of the source-level EEG tensor. Hence, the coupling was first applied between the sensor-level EEG and fMRI tensors (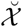 and *𝒴*) and then the resultant common temporal and participant loading matrices were <u>used as a known prior for</u> PARAFAC on source-level EEG 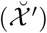. The spectral loading matrix of the sensor-level EEG were used for the initialization of the PARAFAC on source-level EEG. The block diagram of the proposed method is shown in Fig. 1.

**Fig. 1:**
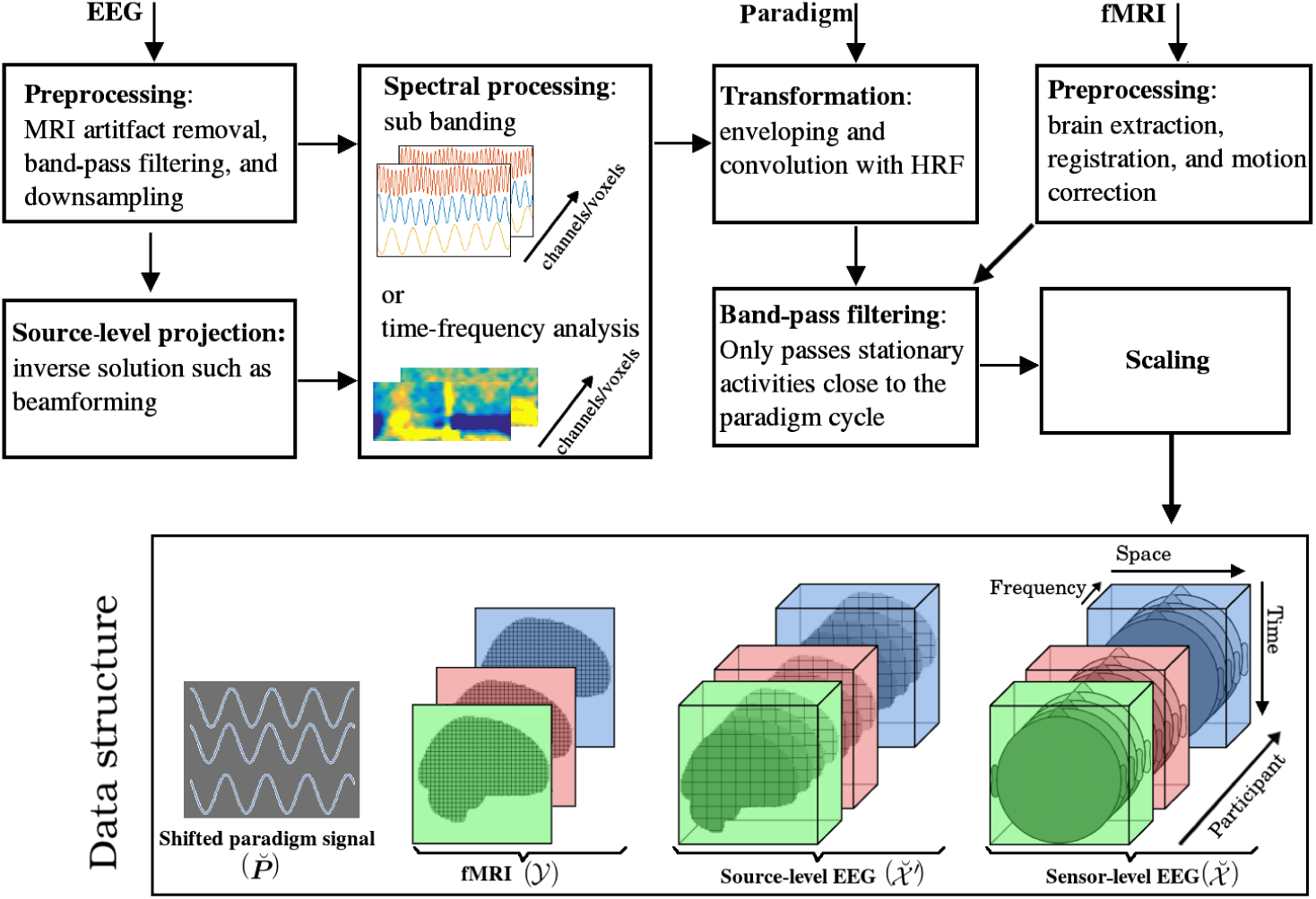
The block diagram illustrates the spatial, temporal, and spectral operations required to create the 4th order EEG and 3rd order fMRI tensors. The EEG and fMRI tensors could be coupled in temporal and participant domains. The paradigm signal could be used as a temporal constraint for the coupled tensor-tensor decomposition.

### 2.4 Data acquisition

Part of the data for this study was obtained from a recent study (Jonmohamadi et al. 2018) and the other part was recorded recently. Participants performed an N-Back working-memory task with alternating 0-Back and 2-Back conditions. During the 0-Back condition, participants responded to the current arrow on the display screen, whereas during the 2-Back condition, participants responded to the arrow two trials earlier. Each arrow was on-screen for 0.5 s. The experimental conditions alternated in a 30 s boxcar with each of the conditions repeated nine times totaling in a 540 s of duration. Before the recording, participants practiced the task at least two times outside the scanner, with at least 80% accuracy achieved.

<u>In total there were 6 participants: 3 males (participants 1, 2, and 3) at the ages of 20, 33, and 38 and 3 females (participants 4, 5, and 6) at the ages of 24, 26, and 22). All participants scored more than 95% correct for the memory task.</u>

EEG data were continuously recorded from 64 channels using the Standard BrainCap MR and BrainAmp MR Plus amplifiers (Brain Products, Munich, Germany) inside an MRI machine. The EEG was acquired using the manufacturer standard cap layout, with the ground electrode located at AFz, reference electrode at FCz, and a drop-down electrode attached centrally to the participants back for the recording of electrocardiography. The impedance of the electrodes were below 10 kOhm.

In order to compute the participant’s individual leadfield, EEG-MR co-registration was achieved by placing Vitamin E capsules, at electrode positions Cz, F5, CP5, and FC6.

MR images were acquired on a 3 T Siemens Skyra, Erlangen, Germany, with a 20-channel head coil. BOLD fMRI data were acquired using a T2*-weighted echo planar imaging (voxel size 3 × 3 × 3 mm).

### 2.5 Data preprocessing

The gradient artefact was removed using realignment parameter-informed artefact correction (Moosmann et al. 2009), and the ballistocardiogram artefacts were rejected using the statistical feature extraction for artifact removal (Liu et al. 2012). EEG was band-passed filtered to 1-20 Hz using a 4th order Butterworth filter, downsampled to 100 Hz, and re-referenced to the FCz electrode. The EEG was then projected into the source-level using the minimum-variance beamformer. The leadfields of the beamformer were calculated using a five-layer realistic finite element model of the head, obtained using the individual MRI structural scan of the participants. The grid size for the beamformer was 8 mm.

It is well known that EEG sources with theta and alpha band activities are associated with working-memory tasks (Debener et al. 2005; Khader et al. 2010; Dong et al. 2015; Stoki et al. 2015; Esposito et al. 2009), therefore, both sensor and source-level EEGs were band-pass filtered to theta and alpha bands. Next, the EEGs were Hilbert enveloped, convolved with a canonical HRF (Glover 1999), and downsampled to the frequency of the fMRI recording (0.4545 Hz). Similarly, the paradigm signal was also convolved with the canonical HRF.

The fMRI processing included brain extraction, registration, and motion correction using the FSL toolbox (Jenkinson et al. 2012) and normalization of the data.

### 2.6 Spatial standardization

In order to present the source-level EEG of different participants in the same space, an 8 mm scanning grid template was created using the T1 brain and warped into the individual brain spaces (MRIs). In the case of fMRI, a 3 mm T1 MNI brain template was used to co-register the individual fMRIs. The white matter was masked out.

### 2.7 Temporal normalization and centring

Centering (removal of the non-zero mean from the data) is required prior to scaling (Bro 1997). Since the task includes 9 cycles of 30 s of 0-Back memory task and 30 s 2-Back memory task, the paradigm signal has the frequency of 1/60 Hz. Centering is achieved by band-pass filtering of the fMRI and EEG time courses of Equation 8 at frequencies of 1/50 to 1/100 Hz. This removes the transient activities and only the stationary ones remain. In order to scale the sensor-level EEG, source-level EEG, and fMRI time courses, the mean of the Frobenius norms of all the time courses of each tensor data were calculated and then each time course divided by the corresponding mean. Using this approach, the sensor and source-level EEG and fMRI time courses have similar variances.

## 3 Results

In order to identify the task related components, a correlation test was performed with the temporal signature of the components and the 4.4 s, 0.0 s, and −4.4 s shifted paradigm signal. Out of the 9 coupled components, components 2-8 had a high correlation coefficient (*>*0.85). The result of the correlation test is shown in Fig. 2, where the subfigure (a) refers to the EEG and subfigure (b) refers to the fMRI components. The first 9 scores are the same between the EEG and fMRI as they were coupled. Besides the coupled components, the component 12 and 18 of the EEG has also higt correlation scores.

**Fig. 2:**
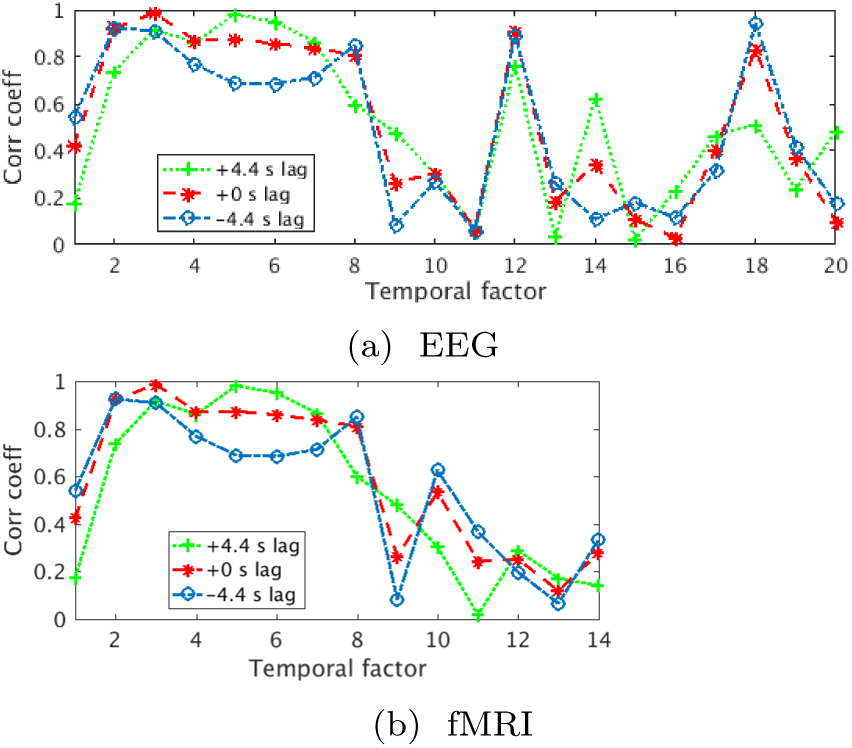
The plot of the correlation coefficients obtained by correlation test between the temporal loading matrix of EEG (upper) and fMRI (lower) with three shifted paradigm signals of +4.4 s, 0.0 s, and −4.4 s. The first 9 factors are coupled between the EEG and fMRI and therefore are the same.

The temporal, participant, spectral, and spatial signatures of the these components are shown in Fig. 3 and 4. According Fig. 2(a), component 2 is in the theta band (5.5 Hz) and several participants, with different extents, have contributed to this component. The EEG sensor map of the coupled component 2 shows the medial frontal channel AFz and surrounding channels pick up highest amount of the memory related theta activity. There are also smaller amount of negativity which is bilateral on the mid posterior channels. The presence of medial frontal theta during memory tasks has been shown in many previous memory studies (Debener et al. 2005; Dong et al. 2015; Michels et al. 2010; Berger et al. 2014) and the slight negativity of the posterior channels in theta band is shown in (Berger et al. 2014) which is consistent with the scalp map shown in Fig. 2(a). The localization of the theta sources has also been reported to have various anterior and posterior origins (Berger et al. 2014) due to the use of different memory tasks. The corresponding EEG source map of the coupled component 2 indicates the origin of theta are being areas which cover frontal cingulate gyrus. According to the fMRI map of the coupled component 2, areas such as premotor, left/medial frontal pole, dorsal cingulate and posterior parietal, shown in Fig. 3(a) in red colour, have increases in BOLD due to the 2-Back memory task. These areas are shown to be the part of the dorsal attention network (Moosmann et al. 2009) and <u>is similar to the maps found by other studies</u> (Owen et al. 2005; Jonmohamadi et al. 2018).

**Fig. 3:**
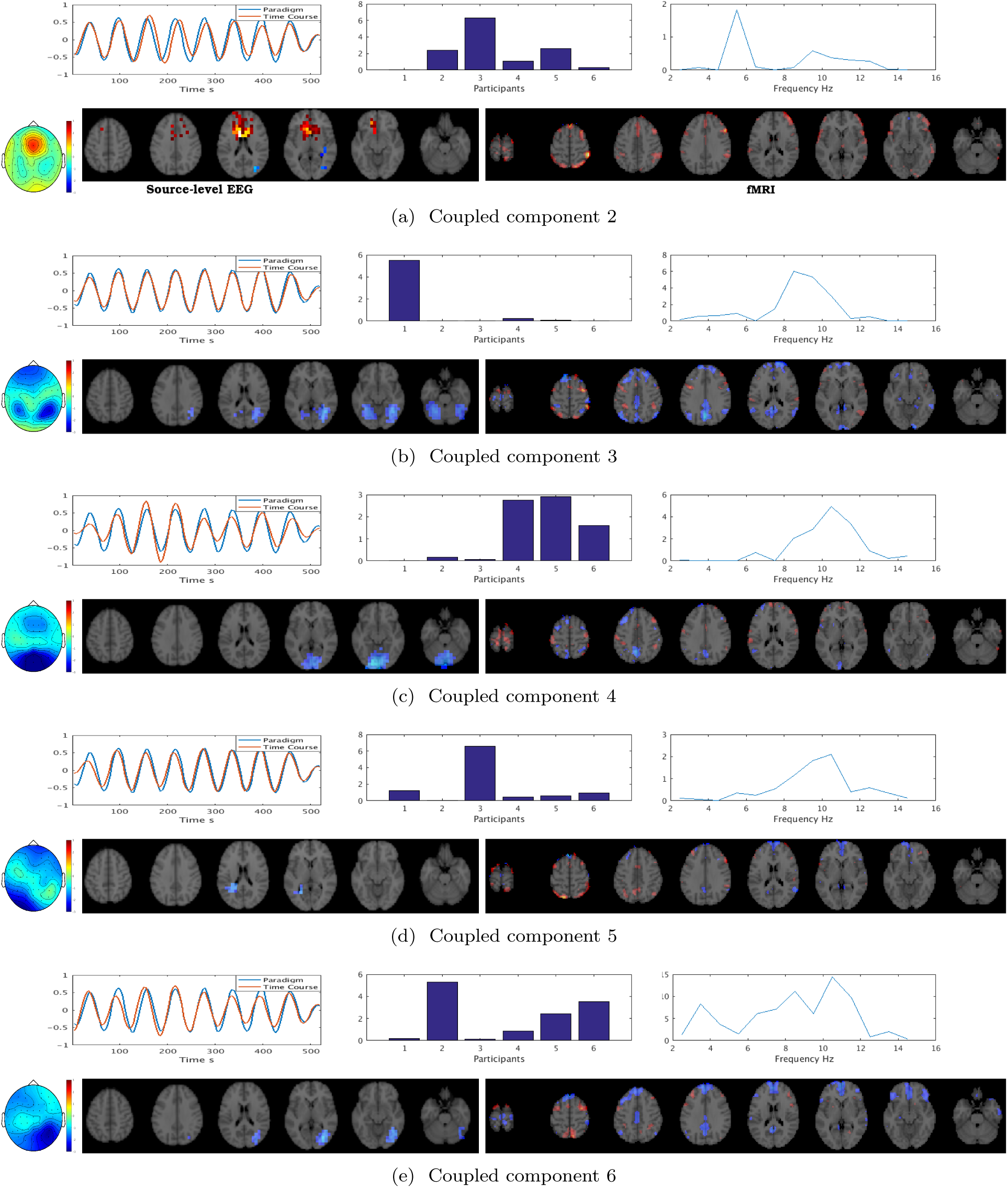
The result of the coupled partial tensor-tensor PARAFAC decomposition on a 4th order EEG tensor and 3rd order fMRI tensor, partially coupled in the temporal and participant domains. In each subfigure, the upper row, from left to right, shows the temporal, participant, and spectral signatures, whereas the lower row, left to right, shows the sensor-level EEG, source-level EEG, and the fMRI signatures of a coupled component. The hot color of the EEG and fMRI maps indicates the positive relation and cold color indicate the negative relation to the paradigm signal. The plot of the temporal signature shows the paradigm signal (in blue) as well as the actual temporal signature of each component (in red). This figure only shows coupled components 2-6. Coupled components 7 and 8 together with the EEG components 12 and 18 are shown in Fig. 4.

**Fig. 4:**
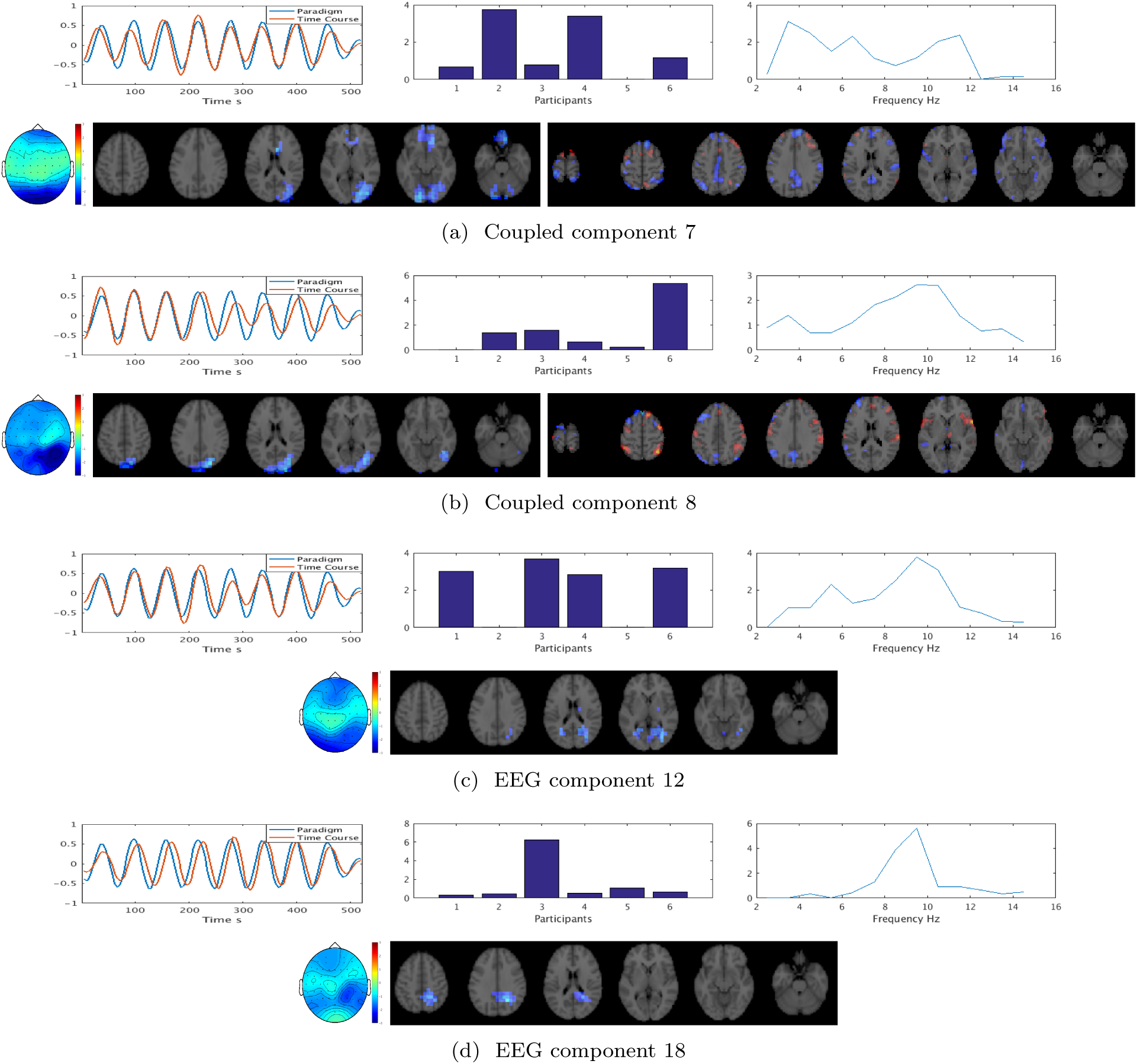
Continue of the Fig. 3: sub figures (a) and (b) are the result of the coupled partial tensor-tensor PARAFAC decomposition on a 4th order EEG tensor and 3rd order fMRI tensor, partially coupled in the temporal and participant domains. In each subfigure, the upper row, from left to right, shows the temporal, participant, and spectral signatures, whereas the lower row, left to right, shows the sensor-level EEG, source-level EEG, and the fMRI signatures of a coupled component. The hot color of the EEG and fMRI maps indicates the positive relation and cold color indicate the negative relation to the paradigm signal. The plot of the temporal signature shows the paradigm signal (in blue) as well as the actual temporal signature of each component (in red). Subfigures (a) and (d) correspond to the components 12 and 18 and are specific to the EEG as they are no coupled to the fMRI.

While there was only one theta band EEG activity related to the memory task, there were several alpha band components which were negatively time locked to the paradigm, either dominated by one participant such as shown Fig. 3(b), (d) and Fig. 4(d) or common with several participants with relatively similar contributions such as Fig. 3(c), (e) and Fig. 4(c). The corresponding fMRI maps of these components are mostly <u>showing the default mode network</u> which include frontal gyrus, medial prefrontal cortex, amygdala, cerebral cortex, and areas of precuneus cortex and posterior cingulate cortex. The scalps maps of these alpha band activities resembles the resting EEG maps as shown in (Ma et al. 2015) where also inter and intra subject differences were acknowledged in the resting state alpha band scalp maps. The EEG source maps show areas such as lateral occipital cortex, occipital pole, and occipital fusiform gyrus as the origins of the alpha activity. <u>The coupled component 4 belongs to the female participants.</u>

According to the temporal signatures, most of the components had similar latencies to the canonical HRF (6 s) but there were two components which maintained consistent lags. Coupled component 5 in Fig. 3(d) remained 2.2 s ahead of the paradigm signal, whereas the component 18 of the EEG in Fig. 4(d) remained 6 s behind the paradigm signal. Both of the mentioned components are dominated by participant 3, with component 5 being from the left occipital area and the component 18 from the right side.

## 4 Discussion

In the field of biomedical imaging, use of more than one imaging modality to capture the variability of a physiological process is a common practice. A primary goal of multimodal data processing is to find components in each of the modalities which are related to the physiological process of interest.

Traditionally, matrix factorization techniques such as PCA and ICA have been used for the purpose of the multimodal data processing. These techniques, rely on the nonphysiological assumption such as orthogonality and statistical independence to achieve uniqueness among the components. In neuroimaging, typical modes of variability in the data are time, space, participant gender, participant age, participant condition, and the tasks in the paradigm. Additionally, mathematical operation on the data such as time-frequency analysis, spatial and temporal filtering can increase the dimension of the data. Matrices by default represent variation of the data in two modes, hence in order to present extra modes of variability in a matrix, unfolding is necessary, which is know to leads the loss of information (Cong et al. 2015). On the other hand, tensors conveniently present multidimensional data and tensor decomposition techniques such as PARAFAC or Tucker decomposition do not impose constraints in the optimization process. Tensor-based analysis of concurrent EEG-fMRI have received increasing attention in recent years (Vanderperren et al. 2010; Karahan et al. 2015; Ferdowsi et al. 2015; Hunyadi et al. 2016, 2017; Acar et al. 2017a,b; Deshpande et al. 2017; Sen and Parhi 2017; Van Eyndhoven et al. 2017; Chatzichristos et al. 2018; Kinney-Lang et al. 2019). However, all the tensor-based fusion of the EEG-fMRI methods, except (Chatzichristos et al. 2018), has been in fact under the matrix-tensor factorization framework, i.e., fMRI in a matrix and EEG in a 3rd order tensor, and only one mode of variability such as participant or time have been used as the common loading vectors for the decomposition of the two modalities. Here, we introduced a framework in which a 4th order EEG tensor (time×space×frequency×participant) was partially coupled in time and subject domains with a 3rd order (time×space×participant) fMRI tensor. Moreover, a matrix containing the paradigm signal and its shifted versions (×9) were used for the initialization of the coupled temporal loading matrix. Out of the nine coupled components, seven components were found to be time-locked to the paradigm signals. The corresponding spatial, temporal, spectral, and participant signatures of these components recapitulated the know well-known resting state and attention networks, available in the literature. Another further two EEG components were also related to the task but were among the non coupled components. <u>One of these two components had a substantial delay of 6 s with respect to the paradigm signal. More over, one coupled component had only contribution from the female participants.</u>

Although the use of the 6 healthy subjects did not show any major finding in regards to the resting state and attention task EEG and fMRI data, but the approach described here demonstrated that using a single process of CTTD, the temporal, spectral, and spatial similarities and differences between the participants can be identified all at once.

## 5 Conclusion

Compared to matrix decomposition, the tensor decomposition techniques are superior due to being able to inherently represent the multidimensional data and the achieving uniqueness without imposing strong assumptions. In recent years the tensor decomposition techniques have gained popularity in the fusion of the biomedical data. Here, a novel tensor-based fusion method was introduced for extraction of the task related EEG and fMRI features. In the proposed method, a single 4th order EEG tensor was partially coupled in time and participant modes to a 3rd order fMRI tensor. The application of the methods was demonstrated on simultaneous EEG-fMRI data from 6 subjects performing 0-Back and 2-Back memory tasks.

## References

Acar E, Lawaetz AJ, Rasmussen MA, Bro R (2013) Structure-revealing data fusion model with applications in metabolomics. In: Engineering in Medicine and Biology Society (EMBC), 2013 35th Annual International Conference of the IEEE, IEEE, pp 6023–6026

Acar E, Papalexakis EE, Grdeniz G, Rasmussen MA, Lawaetz AJ, Nilsson M, Bro R (2014) Structure-revealing data fusion. BMC bioinformatics 15(1):239

Acar E, Levin-Schwartz Y, Calhoun V, Adal T (2017a) ACMTF for fusion of multi-modal neuroimaging data and identification of biomarkers,. In: 25th European Signal Processing Conf.(EUSIPCO-2017)

Acar E, Levin-Schwartz Y, Calhoun VD, Adali T (2017b) Tensor-based fusion of EEG and fMRI to understand neurological changes in schizophrenia. In: 2017 IEEE International Symposium on Circuits and Systems (ISCAS), IEEE, pp 1–4

Andersson CA, Bro R (2000) The N-way toolbox for MATLAB. Chemometrics and intelligent laboratory systems 52(1):1–4

Babiloni C, Babiloni F, Carducci F, Cappa S, Cincotti F, Del Percio C, Miniussi C, Moretti DV, Pasqualetti P, Rossi S (2004) Human cortical EEG rhythms during longterm episodic memory task. A high-resolution EEG study of the HERA model. Neuroimage 21(4):1576–1584

Berger B, Omer S, Minarik T, Sterr A, Sauseng P (2014) Interacting memory systems–does EEG alpha activity respond to semantic long-term memory access in a working memory task? Biology 4(1):1–16

Bro R (1997) PARAFAC.

Tutorial and applications. Chemometrics and intelligent laboratory systems 38(2):149–171

Calhoun VD, Adali T (2012) Multisubject independent component analysis of fMRI: a decade of intrinsic networks, default mode, and neurodiagnostic discovery. IEEE reviews in biomedical engineering 5:60–73

Calhoun VD, Adal T, Kiehl KA, Astur R, Pekar JJ, Pearlson GD (2006) A method for multitask fMRI data fusion applied to schizophrenia. Human brain mapping 27(7):598–610

Calhoun VD, Liu J, Adal T (2009) A review of group ICA for fMRI data and ICA for joint inference of imaging, genetic, and ERP data. Neuroimage 45(1):S163–S172

Carroll JD, Chang JJ (1970) Analysis of individual differences in multidimensional scaling via an N-way generalization of Eckart-Young decomposition. Psychometrika 35(3):283–319

Chatzichristos C, Davies M, Escudero J, Kofidis E, Theodoridis S (2018) Fusion of EEG and fMRI via Soft Coupled Tensor Decompositions. In: 2018 26th European Signal Processing Conference (EUSIPCO), IEEE, pp 56–60

Cichocki A, Mandic D, De Lathauwer L, Zhou G, Zhao Q, Caiafa C, Phan HA (2015) Tensor decompositions for signal processing applications: From two-way to multiway component analysis. IEEE Signal Processing Magazine 32(2):145–163

Cong F, He Z, Hmlinen J, Leppnen PH, Lyytinen H, Cichocki A, Ristaniemi T (2013) Validating rationale of group-level component analysis based on estimating number of sources in EEG through model order selection. Journal of neuroscience methods 212(1):165–172

Cong F, Lin QH, Kuang LD, Gong XF, Astikainen P, Ristaniemi T (2015) Tensor decomposition of EEG signals: a brief review. Journal of neuroscience methods 248:59–69

Debener S, Ullsperger M, Siegel M, Fiehler K, Von Cramon DY, Engel AK (2005) Trial-by-trial coupling of concurrent electroencephalogram and functional magnetic resonance imaging identifies the dynamics of performance monitoring. Journal of Neuroscience 25(50):11730–11737

Delorme A, Makeig S (2004) EEGLAB: an open source toolbox for analysis of single-trial EEG dynamics including independent component analysis. Journal of neuroscience methods 134(1):9–21

Deshpande G, Rangaprakash D, Oeding L, Cichocki A, Hu XP (2017) A new generation of brain-computer interfaces driven by discovery of latent EEG-fMRI linkages using tensor decomposition. Frontiers in neuroscience 11:246

Dien J (2012) Applying principal components analysis to event-related potentials: a tutorial. Developmental neuropsychology 37(6):497–517

Dong S, Reder LM, Yao Y, Liu Y, Chen F (2015) Individual differences in working memory capacity are reflected in different ERP and EEG patterns to task difficulty. Brain research 1616:146–156

Eichele T, Rachakonda S, Brakedal B, Eikeland R, Calhoun VD (2011) EEGIFT: group independent component analysis for event-related EEG data. Computational intelligence and neuroscience 2011:9

Esposito F, Aragri A, Piccoli T, Tedeschi G, Goebel R, Di Salle F (2009) Distributed analysis of simultaneous EEG-fMRI time-series: modeling and interpretation issues. Magnetic resonance imaging 27(8):1120–1130

Farias RC, Cohen JE, Comon P (2016) Exploring multimodal data fusion through joint decompositions with flexible couplings. IEEE Transactions on Signal Processing 64(18):4830–4844

Feige B, Spiegelhalder K, Kiemen A, Bosch OG, van Elst LT, Hennig J, Seifritz E, Riemann D (2017) Distinctive time-lagged resting-state networks revealed by simultaneous EEG-fMRI. Neuroimage 145:1–10

Ferdowsi S, Abolghasemi V, Sanei S (2015) A new informed tensor factorization approach to EEGfMRI fusion. Journal of neuroscience methods 254:27–35

Glover GH (1999) Deconvolution of impulse response in event-related BOLD fMRI1. Neuroimage 9(4):416–429

Harshman RA (1970) Foundations of the PARAFAC procedure: Models and conditions for an” explanatory” multimodal factor analysis

Hunyadi B, Van Paesschen W, De Vos M, Van Huffel S (2016) Fusion of electroencephalography and functional magnetic resonance imaging to explore epileptic network activity. In: Signal Processing Conference (EUSIPCO), 2016 24th European, IEEE, pp 240–244

Hunyadi B, Dupont P, Van Paesschen W, Van Huffel S (2017) Tensor decompositions and data fusion in epileptic electroencephalography and functional magnetic resonance imaging data. Wiley Interdisciplinary Reviews: Data Mining and Knowledge Discovery 7(1):e1197

Jenkinson M, Beckmann CF, Behrens TE, Woolrich MW, Smith SM (2012) Fsl. Neuroimage 62(2):782–790

Jonmohamadi Y, Jones RD (2015) Source-space ICA for MEG source imaging. Journal of neural engineering 13(1):016005

Jonmohamadi Y, Poudel G, Innes C, Jones R (2014a) Source-space ICA for EEG source separation, localization, and time-course reconstruction. NeuroImage 101:720–737

Jonmohamadi Y, Poudel G, Innes C, Weiss D, Krueger R, Jones R (2014b) Comparison of beamformers for EEG source signal reconstruction. Biomedical Signal Processing and Control 14:175–188

Jonmohamadi Y, Forsyth A, McMillan R, Muthukumaraswamy S (2018) Constrained temporal parallel decomposition for EEG-fMRI fusion. Journal of neural engineering

Karahan E, Rojas-Lopez PA, Bringas-Vega ML, Valdes-Hernandez PA, Valdes-Sosa PA (2015) Tensor analysis and fusion of multimodal brain images. arXiv preprint 150606040

Khader PH, Jost K, Ranganath C, Rsler F (2010) Theta and alpha oscillations during working-memory maintenance predict successful long-term memory encoding. Neuroscience letters 468(3):339–343

Kinney-Lang E, Ebied A, Auyeung B, Chin RF, Escudero J (2019) Introducing the Joint EEG-Development Inference (JEDI) Model: A Multi-way, Data Fusion Approach for Estimating Paediatric Developmental Scores Via EEG. IEEE Transactions on Neural Systems and Rehabilitation Engineering

Kronland-Martinet R, Morlet J, Grossmann A (1987) Analysis of sound patterns through wavelet transforms. International Journal of Pattern Recognition and Artificial Intelligence 1(02):273–302

Lagerlund TD, Sharbrough FW, Busacker NE (1997) Spatial filtering of multichannel electroencephalographic recordings through principal component analysis by singular value decomposition. Journal of clinical neurophysiology 14(1):73–82

Liu Z, de Zwart JA, van Gelderen P, Kuo LW, Duyn JH (2012) Statistical feature extraction for artifact removal from concurrent fMRI-EEG recordings. Neuroimage 59(3):2073–2087

Ma L, Minett JW, Blu T, Wang WS (2015) Resting state EEG-based biometrics for individual identification using convolutional neural networks. In: 2015 37th Annual International Conference of the IEEE Engineering in Medicine and Biology Society (EMBC), IEEE, pp 2848–2851

Makeig S, Debener S, Onton J, Delorme A (2004) Mining event-related brain dynamics. Trends in cognitive sciences 8(5):204–210

Martinez-Montes E, Valds-Sosa PA, Miwakeichi F, Goldman RI, Cohen MS (2004) Concurrent EEG/fMRI analysis by multiway partial least squares. NeuroImage 22(3):1023–1034

Michels L, Bucher K, Lchinger R, Klaver P, Martin E, Jeanmonod D, Brandeis D (2010) Simultaneous EEG-fMRI during a working memory task: modulations in low and high frequency bands. PloS one 5(4):e10298

Mijovi B, De Vos M, Vanderperren K, Van Huffel S (2012) Improving spatiotemporal characterisation of cognitive processes with data-driven EEG-fMRI analysis. Prilozi 33(1):373–390

Mitra A, Snyder AZ, Hacker CD, Raichle ME (2014) Lag structure in resting-state fMRI. Journal of neurophysiology 111(11):2374–2391

Miwakeichi F, Martnez-Montes E, Valds-Sosa PA, Nishiyama N, Mizuhara H, Yamaguchi Y (2004) Decomposing EEG data into spacetimefrequency components using parallel factor analysis. NeuroImage 22(3):1035–1045

Moosmann M, Schnfelder VH, Specht K, Scheeringa R, Nordby H, Hugdahl K (2009) Realignment parameter-informed artefact correction for simultaneous EEGfMRI recordings. NeuroImage 45(4):1144–1150

Mrup M, Hansen LK, Herrmann CS, Parnas J, Arnfred SM (2006) Parallel factor analysis as an exploratory tool for wavelet transformed event-related EEG. NeuroImage 29(3):938–947

Ogawa S, Lee TM, Kay AR, Tank DW (1990) Brain magnetic resonance imaging with contrast dependent on blood oxygenation. Proceedings of the National Academy of Sciences 87(24):9868–9872

Owen AM, McMillan KM, Laird AR, Bullmore E (2005) N-back working memory paradigm: A meta-analysis of normative functional neuroimaging studies. Human brain mapping 25(1):46–59

Rivet B, Duda M, Gurin-Dugu A, Jutten C, Comon P (2015) Multimodal approach to estimate the ocular movements during EEG recordings: a coupled tensor factorization method. In: Engineering in Medicine and Biology Society (EMBC), 2015 37th Annual International Conference of the IEEE, IEEE, pp 6983–6986

Seichepine N, Essid S, Fvotte C, Capp O (2014) Soft Nonnegative Matrix Co-Factorization. IEEE Transactions on Signal Processing 62(22):5940–5949

Sen B, Parhi KK (2017) Extraction of common task signals and spatial maps from group fmri using a parafac-based tensor decomposition technique. In: Acoustics, Speech and Signal Processing (ICASSP), 2017 IEEE International Conference on, IEEE, pp 1113–1117

Soong AC, Koles ZJ (1995) Principal-component localization of the sources of the background EEG. IEEE Transactions on Biomedical Engineering 42(1):59–67

Sorber L, Van Barel M, De Lathauwer L (2015) Structured data fusion. IEEE Journal of Selected Topics in Signal Processing 9(4):586–600

Stoki M, Milovanovi D, Ljubisavljevi MR, Nenadovi V, uki M (2015) Memory load effect in auditoryverbal shortterm memory task: EEG fractal and spectral analysis. Experimental brain research 233(10):3023–3038

Valdes-Sosa PA, Sanchez-Bornot JM, Sotero RC, Iturria-Medina Y, Aleman-Gomez Y, Bosch-Bayard J, Carbonell F, Ozaki T (2009) Model driven EEG/fMRI fusion of brain oscillations. Human brain mapping 30(9):2701–2721

Van Eyndhoven S, Hunyadi B, De Lathauwer L, Van Huffel S (2017) Flexible fusion of electroencephalography and functional magnetic resonance imaging: Revealing neuralhemodynamic coupling through structured matrixtensor factorization. In: Signal Processing Conference (EUSIPCO), 2017 25th European, IEEE, pp 26–30

Van Veen BD, Van Drongelen W, Yuchtman M, Suzuki A (1997) Localization of brain electrical activity via linearly constrained minimum variance spatial filtering. IEEE Transactions on biomedical engineering 44(9):867–880

Vanderperren K, De Vos M, Mijovi B, Ramautar JR, Novitskiy N, Vanrumste B, Stiers P, Van den Bergh BRH, Wagemans J, Lagae L (2010) PARAFAC on ERP data from a visual detection task during simultaneous fMRI acquisition. In: Proc. of the International Biosignal Processing Conference., Berlin, Germany, vol 103, pp 1–4

Vervliet N, Debals O, Sorber L, Van Barel M, De Lathauwer L (2016) Tensorlab user guide. Available on: http://www.tensorlab.net

